# Oscillation-based connectivity architecture is dominated by an intrinsic spatial organization, not cognitive state or frequency

**DOI:** 10.1101/2020.07.28.225698

**Authors:** Parham Mostame, Sepideh Sadaghiani

**Affiliations:** Psychology Department, University of Illinois at Urbana-Champaign, Urbana, IL; Beckman Institute for Advanced Science and Technology, University of Illinois at Urbana-Champaign, Urbana, IL

## Abstract

Functional connectivity (FC) of neural oscillations (~1-150Hz) is thought to facilitate neural information exchange across brain areas by forming malleable neural ensembles in the service of cognitive processes. However, neural oscillations and their FC are not restricted to certain cognitive demands and continuously unfold in all cognitive states. To what degree is the spatial organization of oscillation-based FC affected by cognitive state or governed by an intrinsic architecture? And what is the impact of oscillation frequency and FC mode (phase-versus amplitude coupling)? Using ECoG recordings of 18 presurgical patients, we quantified the state-dependency of oscillation-based FC in five canonical frequency bands and across an array of 6 task states. For both phase- and amplitude coupling, static FC analysis revealed a spatially largely state-invariant (i.e. intrinsic) component in all frequency bands. Further, the observed intrinsic FC pattern was spatially similar across all frequency bands. However, temporally independent FC dynamics in each frequency band allow for frequency-specific malleability in information exchange. In conclusion, the spatial organization of oscillation-based FC is largely stable over cognitive states, i.e. primarily intrinsic in nature, and shared across frequency bands. The state-invariance is in line with prior findings at the other temporal extreme of brain activity, the infraslow range (~<0.1Hz) observed in fMRI. Our observations have implications for conceptual frameworks of oscillation-based FC and the analysis of task-related FC changes.

## 1 Introduction

Oscillation-based functional connectivity (FC) is thought to be essential for neuro-cognitive processing (Engel et al. 2013; Singer 1999; Varela et al. 2001). Depending on the particular frequency band, such FC supports performance in the broadest range of cognitive and behavioral domains, from perception and motor output (Khanna and Carmena 2015; VanRullen 2016) to attention (Jensen et al. 2014; S. Palva and Palva 2007; Sadaghiani and Kleinschmidt 2016) and language processing (Rimmele et al. 2018). Oscillation-based FC can be conceptualized in terms of two major modes (Engel et al. 2013; Mostame and Sadaghiani 2020): (1) phase coupling denoting synchronization of the phase of neural activity of distinct brain regions over multiple consecutive oscillation cycles (Lachaux et al. 1999; Nolte et al. 2004), and (2) amplitude coupling denoting synchronization of the magnitude of neural oscillations between regions (Brookes et al. 2011). These FC modes have traditionally been evaluated across a small set of brain regions, and temporally confined to relatively brief task-related processes (Singer, 1999; Uhlhaas et al., 2009). A more recent advance is to derive large-scale connectivity maps from electroencephalography (EEG) or magnetoencephalography (MEG) source space (e.g. Deligianni et al. 2014; Hipp and Siegel 2015; P. Tewarie et al. 2016) and intracranial data (Betzel et al. 2019; Kucyi et al. 2018). This advance is grounded in the understanding that the comprehensive FC organization is of functional importance, at least partially reflecting prior knowledge stored in the connectome (Sadaghiani and Kleinschmidt 2013; Singer 2013).

The degree to which such large-scale FC organization of oscillatory rhythms is modulated by cognitive context is largely unknown. Brain oscillations persist to the same extent in the absence of explicit tasks, i.e. during the so-called resting state. For a century, task-independent ongoing oscillations have been the hallmark of EEG (Berger 1930). The spectral properties of such ongoing activity (“quantitative EEG”) and their cross-areal connectivity during resting state have proven informative for understanding brain function and dysfunction (Kanda et al. 2009; Sadaghiani et al. 2019). The omnipresence and functional importance of electrophysiological oscillations and FC across all cognitive states therefore begs the question of whether they are governed by a state-invariant, i.e. intrinsic, spatial organization.

Several source-localized MEG and EEG studies hint at the presence of a cognitive state-invariant spatial component in oscillation-based FC. These studies have reported the presence of intrinsic connectivity networks and whole-brain FC architecture at resting state (Colclough et al. 2016; Hillebrand et al. 2012; Sockeel et al. 2016; Wirsich et al. 2017). Other studies have confirmed these findings using intracranial recordings of presurgical patients (electrocorticography or ECoG) (Betzel et al. 2019; K. C. Fox et al. 2018; Kucyi et al. 2018) in the absence of the ill-posed problem of EEG/MEG source-reconstruction (J. M. Palva et al. 2018; Schoffelen and Gross 2009). However, these studies have not quantitatively assessed (dis)similarity of oscillation-based FC across different cognitive states.

A few neurophysiological studies have taken important steps towards identifying a state-invariant component in oscillation-based FC across cognitive states. Kramer et al. analyzed daylong ECoG recordings across various levels of arousal and states of consciousness. Static FC across electrodes during periods >100 secs displayed consistent spatial organization over the course of the day. However, this study did not directly and quantitatively contrast FC across the different consciousness levels. A subsequent scalp EEG study identified high spatial correlation of static (>100s) FC organization in sensor space over different sleep stages and uncontrolled wakefulness (r>0.75) (Chu et al. 2012). However, different cognitive activities were not dissociated during the waking period. It is therefore unclear how these findings relate to trial-based oscillatory FC commonly investigated in cognitive neuroscience. A more recent scalp EEG study showed that phase coupling in reconstructed source-space is consistent across tasks (resting state, video viewing, and flashing gratings), with FC clusters that are reproducible across frequency bands (Nentwich et al. 2020). However, while promising the latter two studies must be interpreted with care due to the methodological limitations imposed by EEG recorded over the scalp, which may lead to spurious FC even in source space. Taken together, these findings motivate direct comparison of static FC organization across cognitive states using high fidelity intracranial data.

Dependence on cognitive context has been well quantified in another research field, namely the study of very slow and aperiodic Blood Oxygen Level Dependent (BOLD) changes observed with functional magnetic resonance imaging (fMRI). The brain at resting state displays co-variation in BOLD signals across specific sets of regions, i.e. intrinsic connectivity networks (Beckmann et al. 2005). Importantly, this intrinsic functional architecture is also present during task, suggesting that task-specific changes to the brain’s fMRI-derived FC spatial organization are small (Beckmann et al. 2005; Cole et al. 2014; Gratton et al. 2018; Hearne et al. 2017; Smith et al. 2009). For example, Cole et al. identified a strong spatial correlation (r=~0.9) between the static FC organization of rest and task data.

However, whether these fMRI-based findings can inform about potential state-invariance in oscillation-based FC is questionable, given that fMRI-derived FC is an indirect measure of neural activity based on aperiodic fluctuations of the BOLD signal in the infra-slow range (mainly <0.01Hz) (M. D. Fox and Raichle 2007; Thompson and Fransson 2015). In contrast, FC as measured by electrophysiological methods reflects real-time neural processes driven by cyclic activity. As such, oscillation-based FC is well-positioned to support long-range communication required for cognitive processes that unfold on the rapid timescale of tens to hundreds of milliseconds (Fell and Axmacher 2011; Gruber et al. 2018; Uhlhaas et al. 2009). Given these fundamentally different characteristics of the processes associated with fMRI-based and oscillation-based FC, the degree of context-dependence of the latter requires dedicated investigation.

Another characteristic of fundamental functional importance unique to oscillation-based FC is its aforementioned frequency-dependence, where specific cognitive functions are associated with oscillatory processes in particular frequencies (e.g. Palva and Palva 2007; Rohenkohl, Bosman, and Fries 2018). Based on this functional specificity of particular oscillation bands and their association with different brain regions, should we expect the oscillation-based FC organization in different frequency bands to differ from each other?

In light of the foregoing, we addressed two central questions in the current study: (1) is the spatial organization of fast oscillation-based FC primarily driven by cognitive operations or is it stable across task states?; (2) is this organization dependent on oscillation frequency? Following these two central investigations, we further asked (3) does frequency-invariant FC organization (if observed) reflect a single broadband coupling process or truly multiple temporally independent and frequency-specific processes within a universal spatial organization?; and finally, (4) does the hypothesized stability of static FC organization over cognitive states and frequency bands depend on the specific connectivity mode, i.e., phase- or amplitude coupling? To address these questions while minimizing the impact of volume conduction on FC (J. M. Palva et al. 2018; Schoffelen and Gross 2009), we study ECoG in patients undergoing clinical evaluation for epilepsy surgery. We further replicate major findings using additional FC measures that suppress the impact of potential volume conduction. In summary, we characterized the state- and frequency-dependency of FC as illustrated in 4 possible scenarios in Fig.1.

**Fig. 1.**
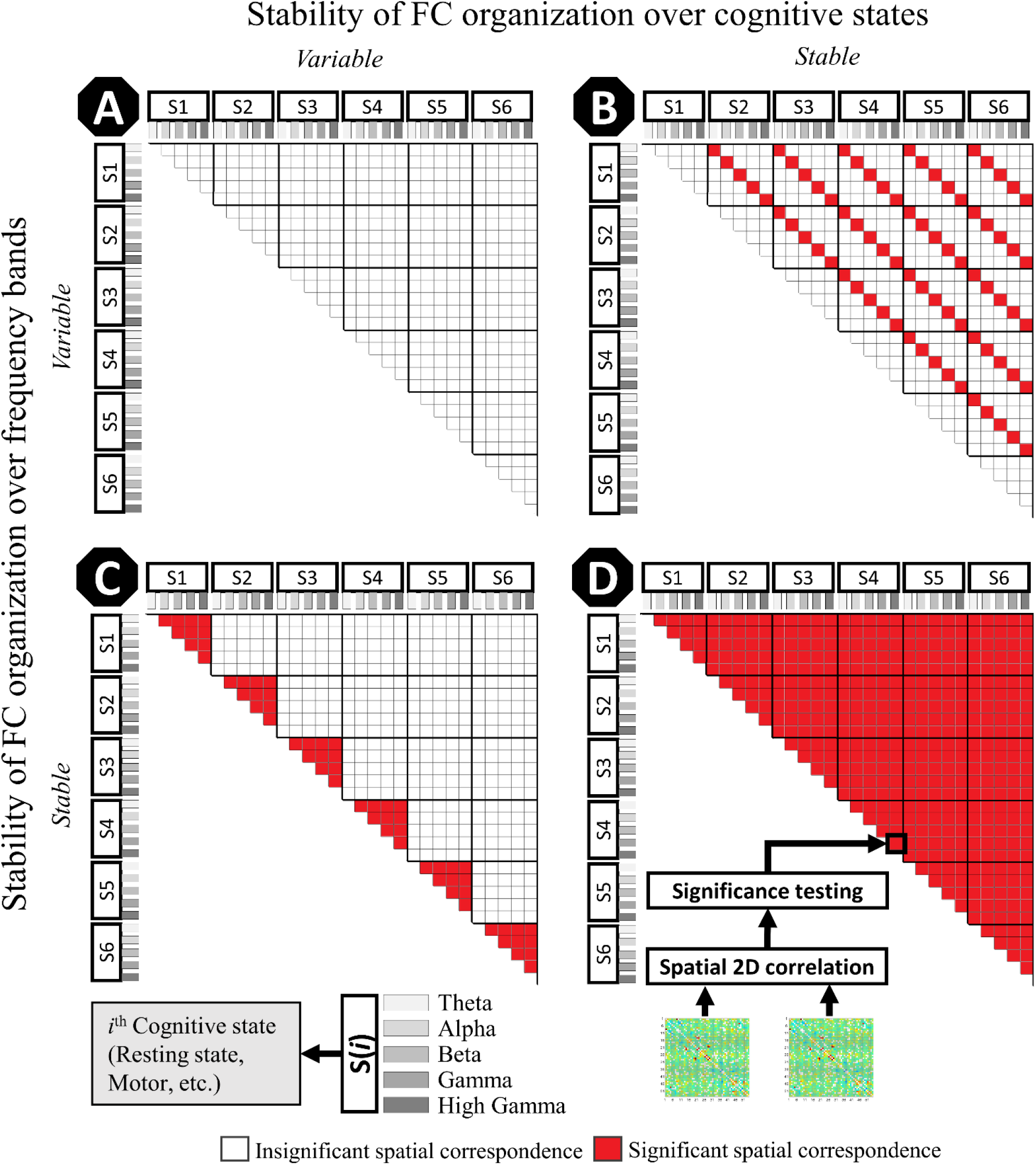
Four hypothetical scenarios regarding stability of the spatial organization of FC across cognitive states and frequency bands. Each cognitive state Mi is subdivided into five frequency bands (theta, alpha, beta, gamma, and high gamma). Each element within the matrix represents the 2D spatial correspondence (quantified as correlation) between corresponding pairs of FC matrices, where red indicates a statistically significant relationship. Our results will be explicitly compared to these four scenarios in Fig. 6.

## 2 Methods

### 2.1 Data

In this study, we used publicly available ECoG recordings during resting state and 3 independent paradigms comprising a total of 6 distinct cognitive states. The data is described in detail in “A Library of Human Electrocorticographic Data and Analyses” Miller (2019) and is available at https://searchworks.stanford.edu/view/zk881ps0522. All patients participated in a purely voluntary manner, after providing informed written consent, under experimental protocols approved by the Institutional Review Board of the University of Washington (#12193). All patient data was anonymized according to IRB protocol, in accordance with HIPAA mandate.

#### 2.1.1 Subjects

From the above-mentioned library, we included only those subjects that had data in at least two paradigms in such a way that maximized the number of subjects for pairs of cognitive states (see below). Cumulatively over the chosen task pairs, the dataset contained 18 subjects. However, not all subjects had data in all of the 6 cognitive states (median number of states across subjects = 3). Subjects - disregarding 3 with missing age information-were on average 30.8 years old (+/−std= 9.3). Average number of electrodes per subject was 53.5 (+/−std=12.7) with an inter-electrode distance of 1cm. All data were sampled at 1000Hz, with a built-in 0.15-200 Hz bandpass filter. Beyond the initial data cleaning performed by Miller et al, portions of the data with excessive inter-ictal activity were removed if necessary.

#### 2.1.2 Behavioral paradigms

In order to increase the statistical power of cross-state comparisons of FC, we used those paradigms that had the highest number of subjects with highest possible number of independent cognitive states (see Table. S2): resting state, cue-based motor task, verb generation task, and 2-back task datasets. These tasks represent a highly diverse set of cognitive states in terms of cognitive demands. Motor and speech tasks have a pre-stimulus interval while the 2-back task employs a continuous block design. As a result, we investigated the intrinsic FC across 6 cognitive states including: resting state (*Rest*), pre-stimulus motor (*Base*_*Motor*_), pre-stimulus speech (*Base*_*Speech*_), post-stimulus motor (*Motor*), post-stimulus speech (*Speech*), and *2Back*.

Resting state (*Rest*): All subjects underwent resting state recordings while fixating on an ‘‘X’’ on the wall 3 m away, for 2-3 minutes. For more information regarding this data, see (Miller et al. 2009, 2012).

Cue-based motor task (*Motor*): seventeen out of the eighteen subjects had motor task data. Patients were instructed to repetitively flex and extend all fingers based on a visual word cue indicating the body part that should be moved (an alternative cue in a separate run instructed tongue movements not analyzed in this study). Movements were self-paced with a rate of ~1-2Hz using the hand contralateral to the side of the cortical grid placement. Each movement interval was 3s long, preceded by a rest interval of the same length (blank screen). To maximize independence between trials, we excluded 0.5s from the two tails of the trials, resulting in a [−2.5, 2.5]s interval relative to cue onset. We analyzed two task states comprising the pre-stimulus ([−2.5, 0]s) and post-stimulus ([0, 2.5]s) intervals. There were between 30 and 75 trials of rest and movement per subject. See Miller et al. (2007) for more detailed information of this task. Miller et al. have reported robust task-related power changes in the high frequency or high gamma range in all subjects, implying that the subjects have task-appropriate electrode coverage suitable for our study.

Verb generation task (*Speech*): Five of the 18 subjects had performed the speech task. The speech data of one subject was excluded due to mismatching electrode grids with respect to the other tasks. Written nouns (approximately 2.5 cm high and 8–12 cm wide) were presented on a screen positioned approximately 1 m from the patient, at the bedside. Patients were asked to speak a verb that was semantically related to the noun. For example, if the noun read “ball”, the patient might say “kick”, or if the noun read “bee”, the patient might say “fly”. Between each 1.6-second noun there was a blank-screen 1.6-second interstimulus interval. To avoid overlap between the trials during data analysis, we defined the trials from −1.5 to 1.5s with respect to stimulus onset. We analyzed two task states comprising the pre-stimulus ([−1.5, 0]s) and post-stimulus ([0, 1.5]s) intervals. Further details of the speech task data are provided in a prior investigation using these data (Miller et al. 2011).

2-back task (*2Back*): Four subjects out of the eighteen had performed the 2back task. Among the available task conditions (0back, 1back, and 2back), we used 2back since it requires higher cognitive involvement (e.g. attention and decision making) which warrants larger possible divergence of cognitive state from resting state and the simple motor task. The stimuli comprised pictures of 50 houses 10 of which were 2back targets, each presented for 600ms. Subjects were instructed to flex their finger when a picture was the same as the one presented two trials earlier (i.e. target). For more details see (Miller 2019). Note that because the Nback task consists of a continuous stream of pictures, we considered task engagement to be continuous. Thus, in the FC estimations of 2Back, we analyze this data exactly same as the resting state data after concatenating all respective blocks (see section 2.3.1 & 2.3.2).

### 2.2 Data preprocessing and estimation of functional connectivity

The publicly available data were initially cleaned (Miller 2019). However, visual inspection of the signal timecourses indicated the need for further cleaning in two subjects. Brief periods of epileptiform activity where thus manually marked and removed from the data of those subjects. Line noise was removed using two 60 Hz and 120 Hz 2nd order Butterworth notch filters. To remove low frequency drift and high frequency noise, data were filtered by low-pass (120 Hz cutoff) and high-pass (2 Hz) 4th order Butterworth filters. For task data, stimulus-locked trials were based on the pre- and post-stimulus intervals described above. All data analyses were implemented in MATLAB, using FieldTrip (Oostenveld et al. 2011) and custom codes that can be found here: https://github.com/connectlab/IntrinsicFC_Mostame.

Our main manuscript focuses on phase coupling. Results for amplitude coupling are provided as supplementary material (Fig. S1 to S4). Phase- and amplitude-coupling are directly compared in section 2.3.4 and 3.4 (Fig. 5 and Fig. S4 of supplementary materials). We estimated FC across five canonical frequency bands including: theta (5-7 Hz), alpha (8-13 Hz), beta (14-30 Hz), gamma (31-60 Hz), and high gamma (61-110 Hz) (Buzsáki and Draguhn 2004). The delta band (~1-4Hz) was not included since the relatively short trial periods didn’t permit its reliable estimation from few oscillation cycles. Below, static and dynamic estimates of phase- and amplitude-coupling are derived.

#### 2.2.1 Static functional connectivity

Static FC was estimated in a single time window for pre-stimulus (*Base*_*Motor*_ *and Base*_*Speech*_) and post-stimulus intervals (*Motor* and *Speech*). Static FC was assessed in terms of phase coupling using the phase locking value (PLV)(Lachaux et al. 1999):

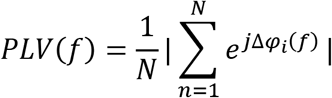

where f is frequency, N is number of trials, and Δφ_*n*_ is the phase difference between the corresponding frequency components of the two electrode signals in trial n.

To derive amplitude coupling, the envelopes of pairwise band-limited signals (estimated through Hilbert transform) were correlated. Subsequently, temporal correlation values of all trials were averaged to obtain a single value for each electrode pair (ranging from −1 to 1), over the corresponding frequency band. To facilitate comparison with PLV, which ranges from 0 to 1, absolute values of amplitude coupling were used.

Static FC for the continuous conditions *Rest* and *2Back* was estimated by averaging the dynamic FC (as defined in section 2.2.2) over time.

#### 2.2.2 Dynamic functional connectivity

Dynamic FC was used to dissociate broad-band versus independent band-limited contributions to the dynamics of the intrinsic FC architecture. Windows were shifted every 1sec, and window length varied as a function of canonical frequency band (75, 100, 200, 400, and 800 cycles for theta, alpha, beta, gamma, and high gamma frequency bands, respectively). Dynamic amplitude coupling was estimated as described in the static framework for each time window. However, dynamic phase coupling was estimated using an alternate method.

In contrast to task-based phase coupling, commonly estimated as phase-lag consistency over trials (Lachaux et al. 1999), *continuous* phase coupling was estimated as phase-lag consistency over time (Mostame and Sadaghiani 2020; Sadaghiani et al. 2012). The above-defined PLV measure adjusted to continuous data is defined as:

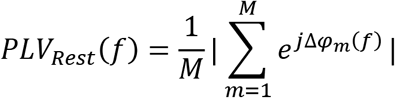

Where M is the number of time samples within the time window.

### 2.3 Intrinsic architecture investigations

#### 2.3.1 Presence of intrinsic architecture

To assess whether FC organization is stable across cognitive states (Fig. 1; scenarios B, D vs. A, C), spatial Pearson correlations were calculated for each state pair. Cross-state correlations were tested against a null model that spatially permuted one of the matrices 500 times (q<0.05; Benjamini-Hochberg FDR corrected for number of subjects × frequencies × task states). Specifically, the phase of the 2D Fourier transform of the matrix was shuffled while keeping the amplitude intact. The odd symmetry of phases was preserved over frequencies in the 2D domain. The matrix was reconstructed using the inverse 2D Fourier transform, and its spatial correlation with the other matrix (which was not shuffled) assessed (Prichard and Theiler 1994; Tewarie et al. 2016; Wirsich et al. 2017).

To determine the effect size of cross-state correlations, spatial stability of FC within *Rest* was estimated for comparison. *Rest* data of each subject was divided into two parts of equal length, and the similarity (2D Pearson correlation) between the static matrices of the two parts was calculated (see Fig. 2).

**Fig. 2.**
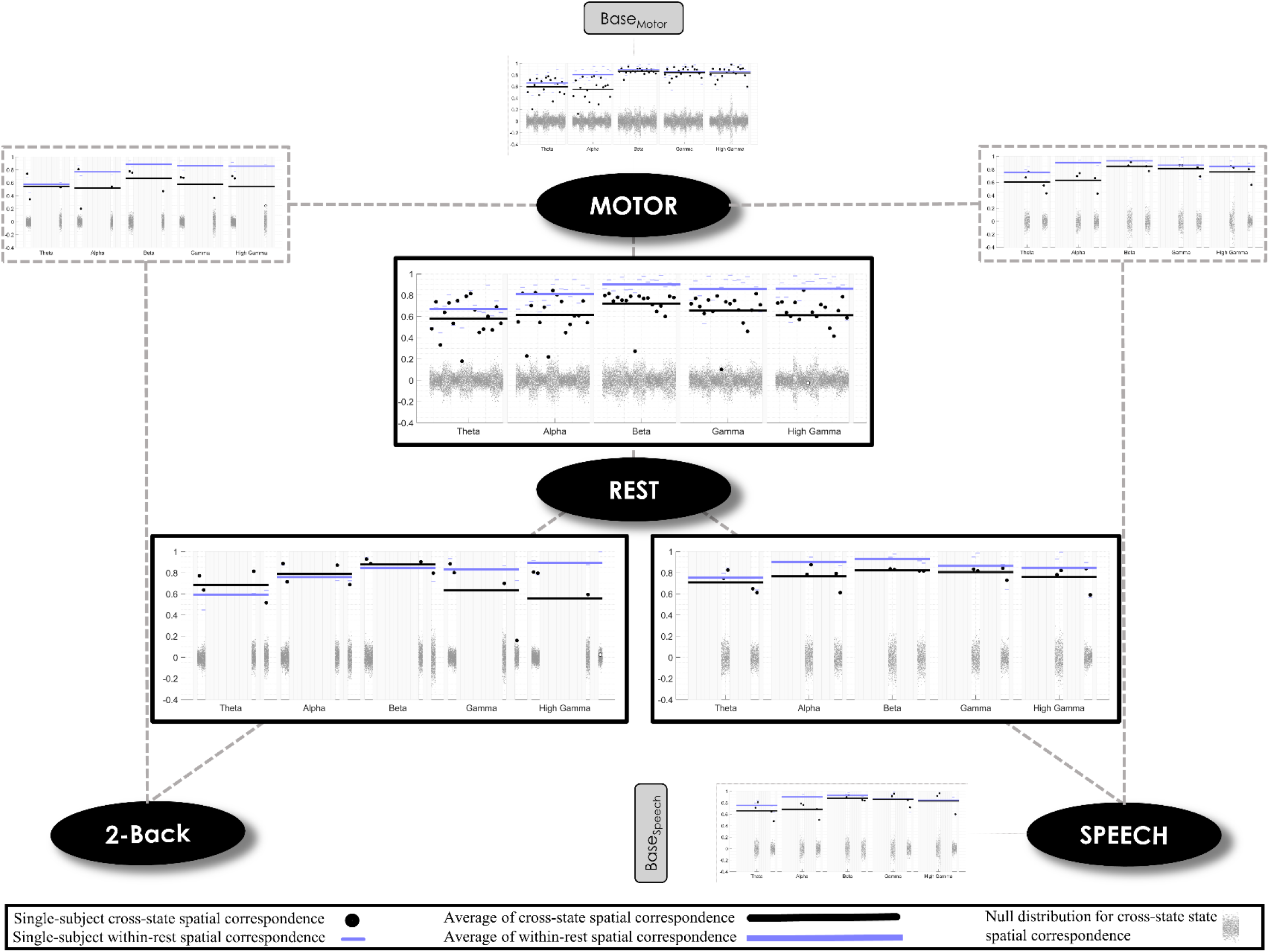
Overview of spatial correspondence between FC matrices of several pairs of cognitive states. States include task-free resting-state (“Rest”), cue-based finger flexing (“Motor”), generating verbs (“Speech”), memory recalling (“2Back”), and the pre-stimulus baseline from the respective tasks (“Base”). Not all subjects had data in all these states. Spatial correlations across any given pair of cognitive states are depicted on the corresponding edge between the state labels. In each subplot, frequency band is on the x-axis and spatial correlation on the y-axis. In each column of the plots, black dots correspond to the cross-state correlation of each subject (black line represents average across subjects). Narrow purple lines represent the within resting-state correlations of FC organization in each subject (purple line shows average across subjects). Gray dot clouds represent the null model for the corresponding subject, frequency band, and cross-state condition. The null model was generated by randomly permuting one of the matrices in 2D Fourier space. Pair-wise correlations significantly exceed the null model in the vast majority of individual subjects and frequency bands (2 exceptions out of 265 pairs), supporting the presence of a stable intrinsic FC organization.

The results for amplitude coupling are provided as supplementary information (Fig. S1).

#### 2.3.2 Shared intrinsic architecture across frequency bands

To assess whether FC in different oscillation frequencies share a similar intrinsic architecture (Fig. 1; scenarios C, D vs. A, B), spatial similarity between the static FC organization of all pairs of frequency bands was calculated. Since the intrinsic architecture is by definition considered stable across cognitive states (Cole et al. 2014; Petersen and Sporns 2015), we estimated such an intrinsic organization for each frequency band by taking the geometrical mean of its static FC organization over all cognitive states (*Rest*, *Base*_*Motor*_, *Base*_*Speech*_, *Motor*, *Speech*, and *2Back*). Then, we calculated spatial correlation of this ‘frequency-specific’ intrinsic architecture over every pair of frequency bands (Fig. 3). Results for amplitude coupling are provided as supplementary information (Fig. S2).

**Fig. 3.**
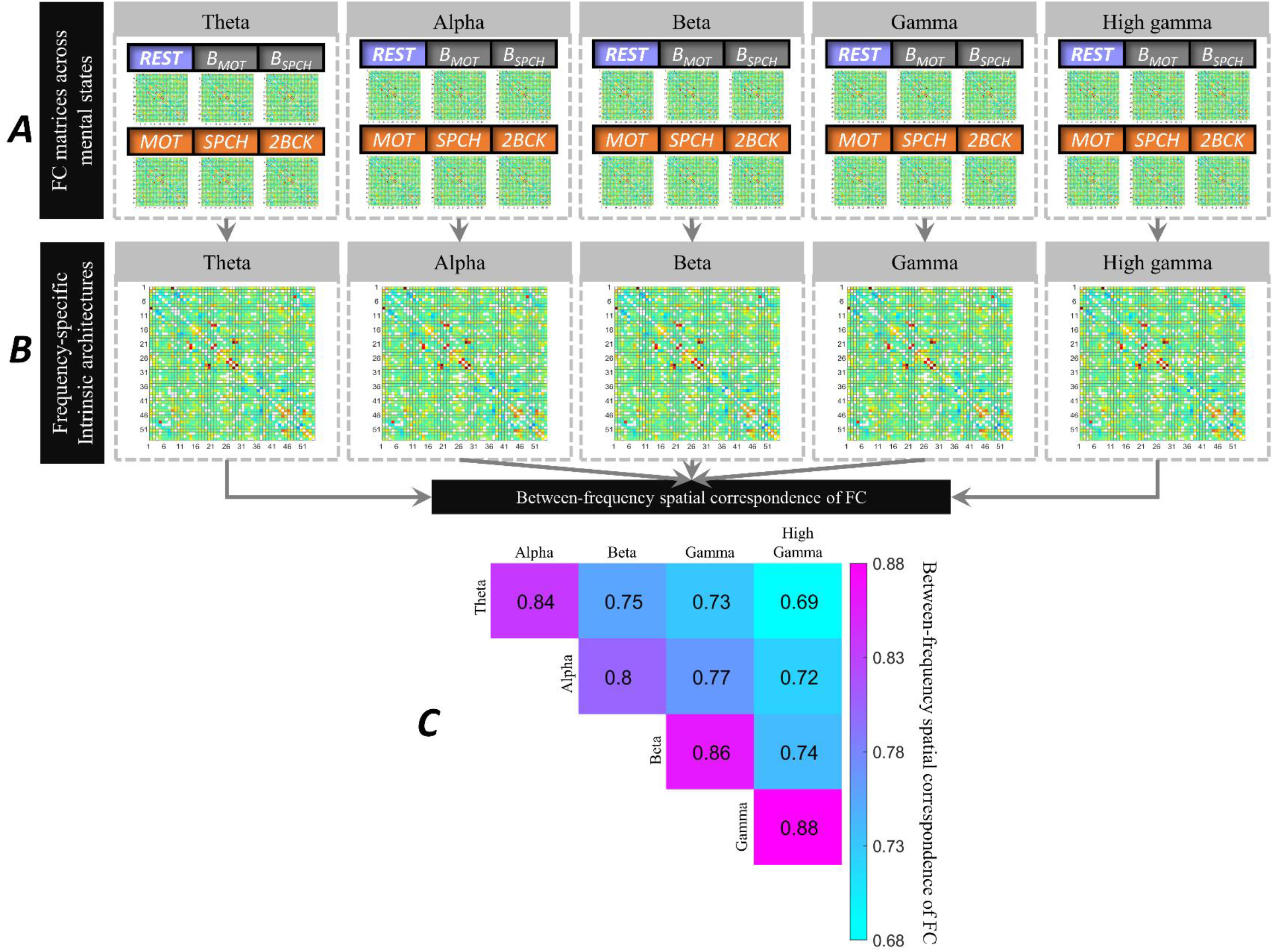
Spatial similarity of intrinsic FC organization across frequency bands. A) State-invariant representations of intrinsic FC organization were calculated separately for each frequency and subject by taking the geometric mean of FC matrices across all cognitive states available for the given subject. B) Then, the ensuing mean intrinsic organization entered spatial correlation analysis across pairs of frequency bands, C) Rows/columns indicate frequency bands from theta to high gamma. The upper triangle indicates degree of correlation. Correlation values of all frequency pairs were statistically significant across all individual subjects. The high correlation values (r>0.69) indicate the presence of an intrinsic organization that is spatially similar across frequency bands.

#### 2.3.3 Broadband vs. band-limited coupling events

Spatial correlation of FC organization across frequency bands may be driven by multiple band-limited processes with a similar spatial organization or, alternatively, a single broadband process. To dissociate between these scenarios, we compared FC timecourses of different frequency bands. The procedure was applied only to the resting state data due to limited length of trial-based task data. To focus on the frequency-invariant connections, only the top 25% strongest connections among all cross-state geometrical mean FC matrices of all bands were considered (mean connectivity strength ± std= 0.60 ± 0.13 pooled over all connections and frequency bands and averaged over subjects). Within the selected connections, temporal correlations between phase coupling timecourses (see 2.2.2) were estimated over pairs of different bands. Finally, for each subject and frequency pair, all temporal correlations were pooled over connections (shown in Fig. 4; corresponding results for amplitude coupling are shown in Fig. S3).

**Fig. 4.**
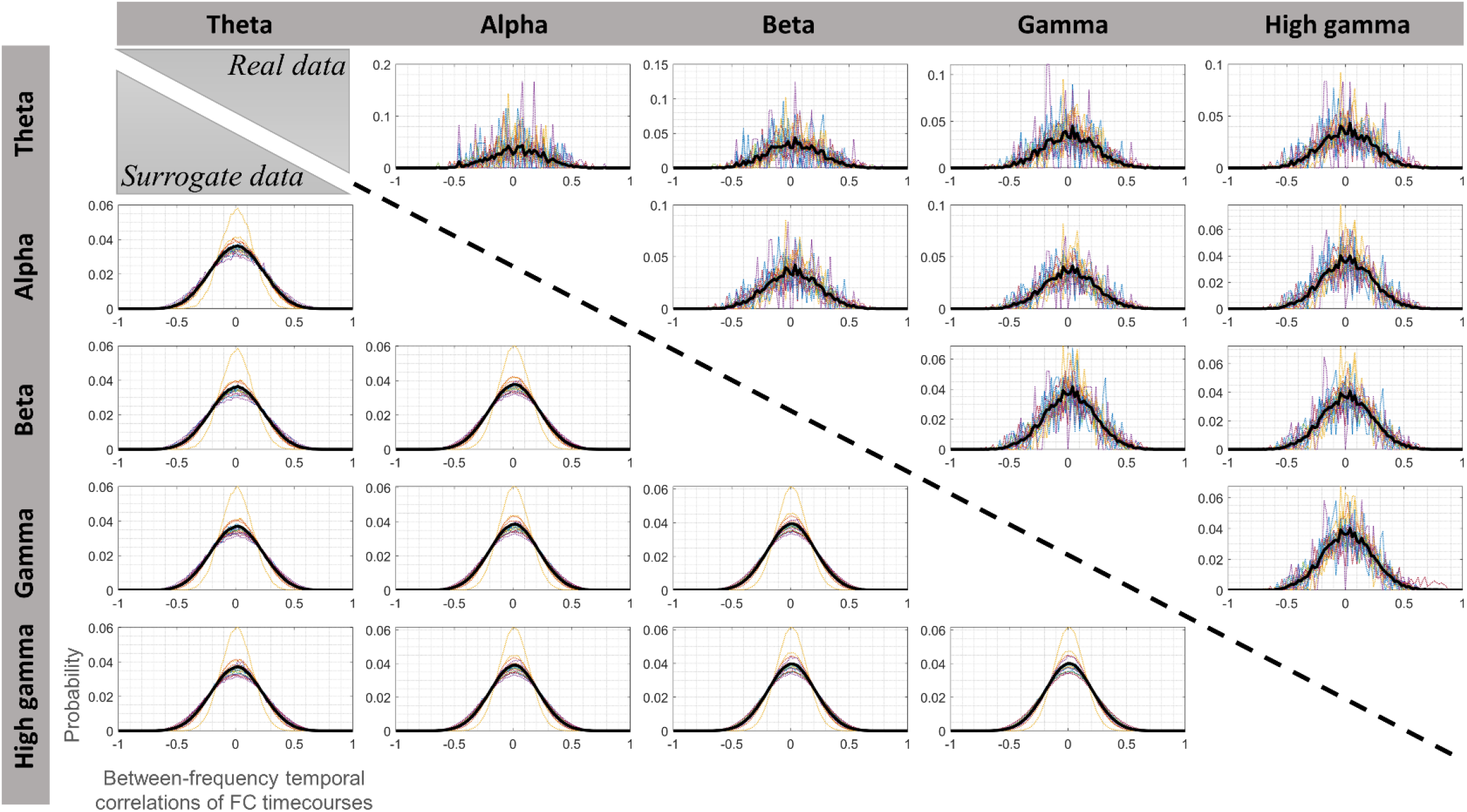
Histogram of temporal correlation values between resting state FC dynamics of different frequency bands for real (upper diagonal) and surrogate (lower diagonal; pooled over 500 repetitions) data. The surrogate data represent temporally phase permuted and thus temporally independent FC time courses. Each panel corresponds to a specific pair of frequency band as labeled. In each panel, histograms averaged over subjects are shown as thick black lines, while single-subject histograms are shown as narrow dotted lines of different colors. The x-axis indicates temporal Pearson correlation values of FC dynamics of the two frequency bands, and the y-axis shows the probability of observing the respective values over electrode pairs. The histograms for real data were zero-centered and symmetric as were those observed for the simulated temporally independent FC dynamics. This observation is consistent with the presence of multiple frequency-specific processes with temporally independent time-varying dynamics rather than a single broadband phenomenon.

To test to what degree the FC dynamics of two bands are systematically correlated, we generated a set of 500 surrogate data by phase-permuting the FC timecourse of one of the frequency bands (for each frequency pair and subject). For each permutation, we extracted the histogram of temporal correlations across all electrode pairs between the original and phase-permuted FC dynamics. The 500 surrogate histograms were pooled into a single histogram for each subject and frequency pair (Fig. 5). To assess if the original histogram was different from the surrogate histogram (indicating systematic correlation between frequency bands in the original data), we compared their mean, std, and skewness (q<0.05; FDR-corrected for all subjects, pairs of frequency bands, and three histogram measures).

**Fig. 5.**
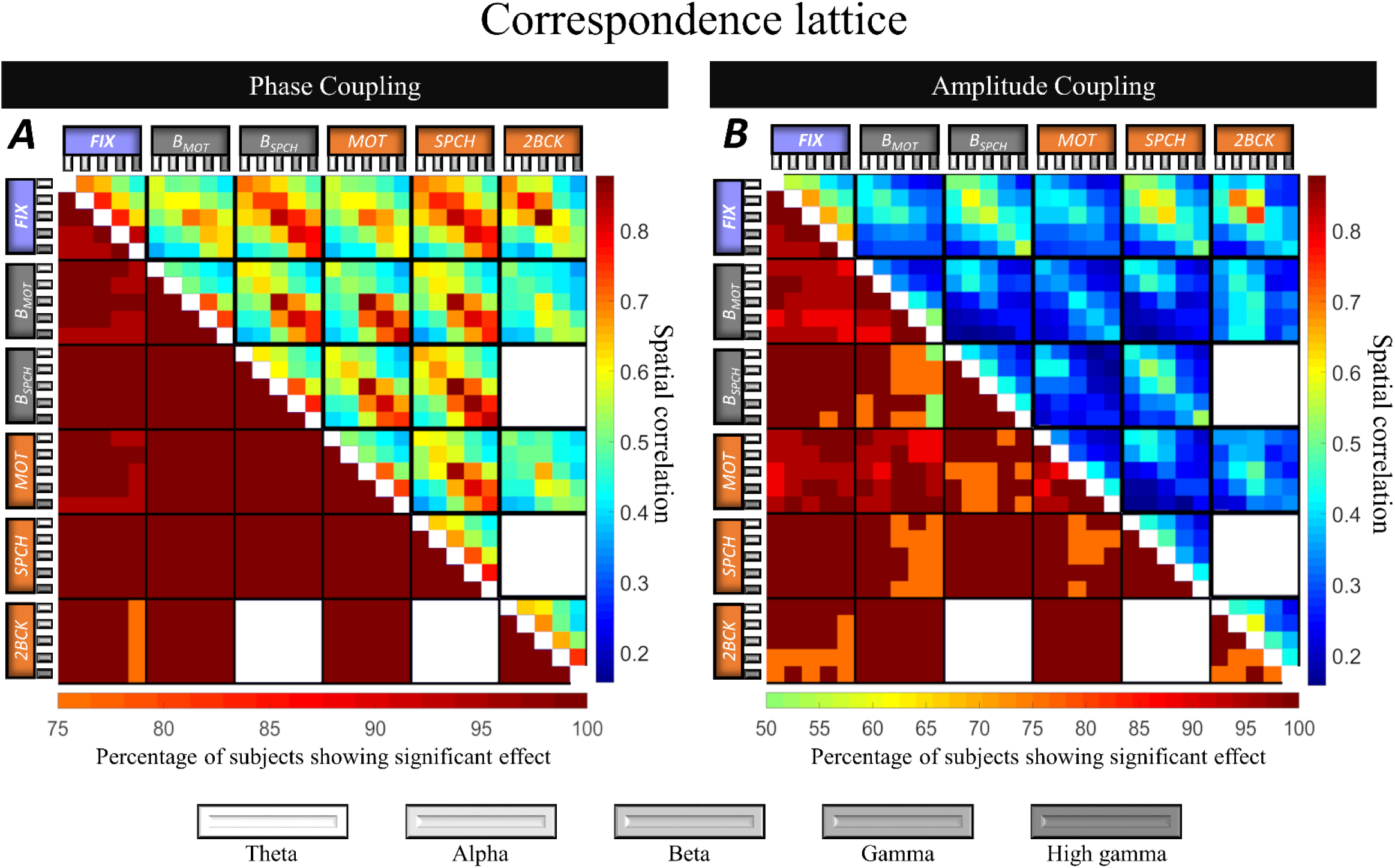
Spatial correlation between FC matrices from all possible pairs of cognitive states and frequency bands (theta to high gamma) for A) phase coupling, and B) amplitude coupling. Upper triangles show correlation values averaged across all subjects, with effect size indicated by color (vertical color bar). For each subject, a set of 500 surrogate data were generated to test the significance of the correlation values. The number of subjects showing significance in each bin of the correspondence lattice is presented in the lower triangles (horizontal color bar). The lower triangles in A and B conform with hypothesis D (Fig. 1). The results suggest that both phase- and amplitude-coupling are governed by an intrinsic FC organization that is consistent across cognitive states and frequency bands.

#### 2.3.4 Shared intrinsic architecture across connectivity measures

To directly assess the spatial similarity of intrinsic FC organization across phase- and amplitude coupling as well as over all frequencies and cognitive states, we estimated spatial Pearson correlation between all possible pairs of FC matrices (across FC measures, cognitive states, and frequency). Finally, we averaged the resulting value of each pair-wise comparison across all subjects (Fig. 5; upper diagonal). We tested the significance of each bin by using the same 2D permutation method described above, phase-permuting one of the two matrices 500 times (q<0.05; FDR corrected by Benjamini-Hochberg method).

### 2.4 Addressing source leakage

Source leakage refers to the simultaneous detection of a particular brain signal (as a source) at several sensors due to volume conduction (Schoffelen and Gross 2009), which may generate spurious connectivity between electrode pairs especially in neurophysiological scalp recordings like EEG and MEG (J. M. Palva et al. 2018). Although much less of a concern for ECoG, we tested whether our results are affected by this potential confound.

First, following equivalent procedures for task and rest conditions as described for PLV, we employed imaginary part of coherency (ImC; introduced by Nolte (2004)). ImC is the most commonly used measure suppressing zero-lag connectivity. This common approach rests on the assumption that electricity spreads quasi-instantaneously, but comes at the cost of also removing real zero-lag connectivity (M. X. Cohen 2015). Therefore, as a second approach, we regressed out electrode distance dependencies from FC values. Given the fact that volume conduction is dependent on electrode distance (Dubey and Ray 2019; Rogers et al. 2019; Rouse et al. 2016), this approach is expected to substantially dampen possible volume conduction effects. Specifically, we fit cubic spline curves to the mean of the FC values within each non-overlapping 1cm range of electrode distance. We subtracted the value of the fitted curve from all corresponding FC values. This procedure removes any collinearity between the two measures resulting from their dependence on electrode distance. Once zero-lag FC or distance dependencies were removed, we estimated the cross-state spatial correlations as in section 2.4.1 (Fig. S5).

## 3 Results

FC matrices were extracted for all subjects, cognitive states, and canonical frequency bands. First, to establish the presence of a cognitive state-invariant, i.e. intrinsic, FC organization, spatial correlation of the FC matrix across different pairs of cognitive states were assessed within each frequency band (section 3.1). Subsequently, the intrinsic FC organization most representative of each frequency was defined as the (geometric) mean of FC matrices across cognitive states in that frequency. Next, this representative organization was spatially compared across all bands to quantify frequency-invariance of FC organization (section 3.2). Then, we compared FC timecourses across frequency bands to address whether the spatial frequency-invariance indeed reflects band-limited coupling in temporally independent frequencies or, alternatively, a broadband coupling phenomenon (section 3.3). Finally, to identify the most likely scenario from the four hypothetical scenarios illustrated in Fig. 1, the spatial similarity across all pairs of FC matrices for all cognitive states and frequency bands was assessed (section 3.4).

### 3.1 A state-invariant intrinsic FC organization

#### 3.1.1 Cross-state spatial correspondence in real vs. surrogate data

Spatial correlations between FC matrices of different cognitive states were assessed in every frequency band (Fig. 2; black data points). All pairs of cognitive states demonstrated strong spatial similarity in all frequencies (average *r* = 0.69; std= 0.18). Spatial correlations were compared to surrogate correlations generated after spatially shuffling one of the two matrices in 2D Fourier space (500 permutations; Fig. 2; Gray dot clouds). At the group level, we compared the subject-specific cross-state correlations (averaged over all cross-state conditions) with the mean value of the corresponding null models (averaged over permutations and all cross-state conditions). For all frequency bands, cross-state spatial similarity exceeded the respective null model (paired *t*-test; all t_17_>3.41; p<0.0017). At the individual level, the spatial similarity across cognitive states exceeded chance level for all individual subjects and frequency bands with only 2 exceptions out of 265 cross-state cases (q<0.05; Benjamini-Hochberg corrected for subjects × frequencies × cognitive states). This outcome indicates the presence of an intrinsic FC organization of phase coupling in all frequency bands. Results for amplitude coupling were comparable (Fig. S1).

#### 3.1.2 Cross-state vs. within-state spatial correspondence

Spatial similarity of FC patterns was also assessed *within* a single cognitive state for comparison. The relatively long duration of resting state recordings allowed comparing time-averaged FC matrices from two equal halves of the run (Fig. 2; purple data points). Overall, spatial similarity *within* resting state was either equivalent or only slightly stronger compared to the similarity *across* cognitive states. Specifically, the average reduction of the *r* values over all cross-state conditions and frequency bands was 10 ± 14% (Mean ± std; see supplementary materials Fig. S2). Therefore, divergence of spatial organization of FC across different cognitive states is comparably small. Results for amplitude coupling are provided in Fig. S1.

### 3.2 Intrinsic FC organization shared across frequency bands

Above, we showed the presence of an intrinsic FC organization that is stable across cognitive states within each of the canonical frequency bands (Fig. 2). Is a unifying state-invariant FC organization shared across frequency bands, or does each specific frequency have its own unique spatial organization? Here, we assess how the state-invariant FC organization of each frequency band spatially correlates with that of other bands. To this end, for each subject and frequency band, we calculated the geometric mean of the FC matrices from all cognitive states as the representative intrinsic organization of that frequency band (Figure 3A). The geometric mean is often used to find the central tendency for different items, emphasizing consistency. In other words, if an electrode pair has a small FC value in even a single cognitive state, that pair obtains a small value in the intrinsic organization matrix since it is not consistent over all cognitive states.

For each subject, we assessed spatial correlations across pairs of representative (mean) intrinsic FC. For each pair of frequencies, we statistically tested the significance of FC similarity using the spatial permutation method described in section 3.1.1 (500 permutations). At the group level, we found significantly higher cross-frequency similarity of FC organization compared to null data (pair-wise *t*-test against the mean correlation from permutations; all *t*_17_>9.37, p<2e-8). At the individual level, the cross-frequency spatial similarity exceeded chance level of the null model for all frequency pairs and all subjects (q<0.05; FDR corrected for 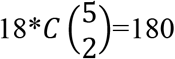 cases of subjects × frequency pairs). Figure 3B shows the spatial correlation values averaged over subjects (upper triangle) as well as the number of subjects showing a significant effect (lower triangle). The strong spatial relationship (*r* = 0.69 to 0.88) across all frequency pairs demonstrates that FC is governed by a universal spatial organization largely shared across frequency bands. Equivalent results from amplitude coupling are provided as supplementary material (Fig. S2).

### 3.3 Temporal dynamics dissociating broadband vs. band-limited FC

Our observation that the intrinsic FC organization is highly similar across frequency bands may be explained by one of two alternative scenarios. Either temporally independent coupling processes in multiple frequency bands indeed share a unifying spatial organization, or a single broadband coupling process (e.g. coupled bursts with sharp on/offset) underlies FC in all frequencies. In the latter case, FC would be temporally correlated over frequencies. To dissociate between the two scenarios, we inspected temporal correlations between connection-wise FC time courses of different frequency bands within the resting state condition because it provides the most data of all cognitive states (2-3 minutes). We only focused on those connections of the intrinsic FC organization that were strong in all frequency bands (top 25% of the representative mean organization discussed in section 2; mean FC= 0.60; std= 0.13; averaged over subjects). This approach ensured that we only included electrode pairs involved in the task-invariant FC organization of all bands. As shown in Fig. 4 (top subplots above the diagonal; ‘real data’), we detected a symmetric histogram of temporal correlation values centered on zero for every subject and pair of frequency bands. The lower subplots below the diagonal of Fig. 4 show an equivalent temporal correlation analysis for surrogate data generated by temporally permuting phases of one of the FC dynamics in Fourier space (500 permutations). The distribution of temporal correlations was highly similar to the corresponding histogram of the real data. Again, similar results were obtained for amplitude coupling (Fig. S3).

To test for an overall tendency towards positive or negative correlations between FC temporal dynamics of each frequency pair, we compared mean, std, and skewness between histograms of the real and null data. The (grand average) difference in the three measures between histograms of the real and null data (averaged across repetitions) were 0.01, 0.01, and 0.02 (±std= 0.03, 0.01, and 0.13), respectively. All differences were negligible in size, suggesting the absence of a broadband process. Moreover, a large proportion of all cases (all subjects, pairs of frequency bands, and the three histogram measures) did not pass the significance test when compared to surrogate data (73, 68, and 95% of all cases for mean, std, and skewness differences, respectively; *q* < 0.05). This observation implies that FC fluctuations in different frequency bands do not temporally coincide (in a linear sense) more frequently than expected by chance. We conclude that the observed spatial concordance of FC organization across frequencies is not generated by a single broadband phenomenon, but rather reflects multiple frequency-specific coupling processes unfolding within the same universal spatial organization.

### 3.4 Stability of spatial organization across phase- and amplitude coupling

In previous sections, we separately tested the stability of FC organization across cognitive states and frequency bands. Next, in order to test the four scenarios presented in Fig. 1, we assessed the stability of FC organization across all possible pairs of cognitive states and frequency bands. We further extended this comprehensive approach to a comparison across different modes of connectivity, i.e. phase- and amplitude coupling.

#### 3.4.1 Spatial organization of phase-coupling

We estimated spatial correlation for all possible pairs of FC matrices across cognitive states and frequencies for each subject separately, resulting in a *correspondence lattice* (Fig. 5). For visualization purposes, the upper triangle of the correspondence lattice in Fig. 5A shows the correspondence lattice averaged over all subjects (*r*= 0.57 ± 0.18). Using the same spatial permutation test as in section 3.1.1, we tested the significance of each bin of the correspondence lattice for each subject. The lower triangle of Fig. 5A reports the proportion of subjects passing significance threshold for each comparison (R=500; q<0.05; Benjamini-Hochberg FDR corrected for all cases of subjects × pairs of cognitive state x frequency bands). We observed evidence for spatial similarity across all pairs of FC matrices in overall 99.3% of the single-subject comparisons, strongly supporting hypothesis D (cf. Fig. 1). This outcome shows that the spatial FC organization is to a large degree consistent across all cognitive states and frequency bands.

#### 3.4.2 Spatial organization of amplitude-coupling

Amplitude coupling is a distinct mode of oscillation-based FC beyond phase, likewise subserving long-range neural communication and cognitive processes (Engel et al. 2013). We have previously hypothesized that FC organization in both modes entails an intrinsic, task-independent component (Mostame and Sadaghiani 2020). Therefore, we asked whether the same observation, i.e. consistency across cognitive states and frequency bands, also holds true for amplitude coupling. We extracted FC matrices of all cognitive states and frequency bands using amplitude coupling as connectivity measure. Fig. 5B visualizes the outcome for amplitude coupling (equivalent to Fig. 5A for phase coupling). Spatial correlation values were moderate to strong (average of upper triangle of 5B *r*= 0.42 ± 0.21). Importantly, as shown in the lower triangle of 5B, spatial correlation of FC matrices across all cognitive states and frequencies were significant in overall 94.7% of the comparisons (q<0.05; Benjamini-Hochberg FDR corrected for subjects × pairs of cognitive state x frequency bands), in line with hypothesis D in Fig. 1. We conclude that the intrinsic architecture is also consistent across cognitive states and frequency bands for amplitude coupling.

#### 3.4.3 Comparing spatial organization between phase- and amplitude-coupling

We observed differences between phase- and amplitude coupling. While spatial correlation values for both were significant compared to their respective null models, values were overall slightly higher for phase coupling (Fig. 5A) than for amplitude coupling (Fig. 5B). Further, as the frequency increased, within-frequency cross-state correlation values increased for phase coupling (diagonals of small squares in Fig. 5A), while they slightly decreased for amplitude coupling (Fig. 5B). This means that task-specific FC changes in phase coupling are more pronounced in lower frequencies (mostly in alpha band), resulting in lower cross-state correlations (Fig. S4; blue bars). By contrast, such task-specific changes are generally stronger in higher frequencies for amplitude coupling (Fig. S4; orange bars). Collectively, these observations suggest the presence of an intrinsic architecture in both phase- and amplitude coupling, but with nuanced differences.

### 3.5 Source leakage contributions

Compared to EEG and MEG, ECoG recordings are much less likely to suffer from volume conduction especially at the scale of ~>1cm inter-electrode distance (Dubey and Ray 2019). Nevertheless, to ascertain that FC stability across cognitive states is largely independent of volume conduction, we replicated our results following suppression of zero-lag connectivity (since electricity spreads quasi-instantaneously) and distance dependence of connectivity (since volume conduction is dependent on physical distance). We used the *Rest*-to-*Motor* comparison due to its large number of subjects (See supplementary section 2 and Fig. S5).

#### 3.5.1 FC measures suppressing zero-lag connectivity (ImC measure)

We found that significant *Rest*-to-*Motor* cross-state FC correspondence persisted in the majority of bands and subjects (63 cases out of 85; with a minimum of 11 out of 17 subjects in each band) even after reducing the data to nonzero-lag connectivity (Fig. S5 B; FDR-corrected at q<0.05). Despite a reduction in effect size likely due to concomitant removal of veridical zero-lag connectivity, the persistence of effects implies that cross-state FC correspondence is not primarily explained by volume conduction effects.

#### 3.5.2 Regressing out distance dependencies from FC

In 82 out of 85 subject-by-frequency cases, spatial organization of FC remained significantly correlated between *Rest* and *Motor* conditions after removing distance dependencies (Fig. S5 C). The large proportion of significant effects even after removal of a substantial part of connectivity indicates that our major findings are not primarily driven by distance dependencies. This persistence suggests that volume conduction contributes no more than a small proportion of the observed consistency of FC organization across cognitive states and frequency bands.

## 4 Discussion

The present study sought to answer whether fast oscillatory coupling was sensitive to cognitive context, or, conversely, associated with a single, state-invariant spatial organization across cognitive domains. We used ECoG signals of presurgical patients during 6 different task states including rest, cue-based motor, speech, 2-Back, and two pre-stimulus intervals corresponding to maintenance of task-set and attention. Across all subjects and all canonical frequency bands, oscillation-based FC spatial organization changed only slightly in response to cognitive state. Moreover, the observed spatial organization was largely similar across all frequency bands. Despite this between-frequency spatial similarity, temporally independent dynamics were detected across frequency bands speaking against a broadband phenomenon. Taken together, our findings suggest that the intrinsic FC of the human brain is characterized by state- and frequency-invariant spatial organization in both phase- and amplitude coupling.

### Stability of oscillation-based FC across cognitive states

Spatial stability of fast oscillation-based FC over cognitive states is in strong agreement with prior observations in fMRI-based FC at infraslow temporal scales. Several neuroimaging studies have shown that the spatial organization of FC remains largely stable across cognitive states (Cole et al. 2014; Gratton et al. 2018; Krienen, Yeo, and Buckner 2014). However, fMRI and electrophysiological measures emphasize partly divergent types of neural processes (Hari and Parkkonen 2015; Hermes et al. 2019; Hermes, Nguyen, and Winawer 2017). fMRI-based FC represents coupling of infraslow and aperiodic fluctuations of activation amplitude. In contrast, coupling of fast oscillatory neural signals likely represent different FC mechanisms. Such strong spatial stability (*r*=0.69; mean over all cross-state comparisons of phase coupling within frequencies) is especially surprising in light of the role of neural oscillations in rapid cognitive processes. It is important to consider the interplay between space and time in FC dynamics. Our observation of spatial stability of FC in various cognitive contexts over the full trial duration does not preclude divergent temporal dynamics in different tasks (see discussion of time-varying dynamics below). Spatial correspondence of FC across states was high but not perfect, suggesting the presence of task-specific adjustments. This latter observation is in line with small but task-specific changes in fMRI-based FC during particular tasks (e.g. Cohen and D’Esposito 2016). Thus, the observed spatial stability is compatible with additional minor but functionally important and task-specific changes in FC patterns.

### Stability of oscillation-based FC across frequency bands

The cognitive state invariance of oscillation-based FC organization was observed in all canonical frequency bands from theta to high gamma, suggesting that this spatial organization is conserved across frequencies. Direct pairwise comparisons of frequency-specific intrinsic FC across frequency bands revealed highly similar FC organization across all bands (*r* = 0.69 to 0.88). This finding is surprising, given that frequency-specific oscillation-based FC has been linked to different cognitive operations, each associated with different sets of brain areas (S. Palva and Palva 2018). For instance, theta, alpha, and gamma band oscillations (both in terms of local power and cross-region coupling) are thought to reflect navigation and memory encoding/retrieval (Backus et al. 2016), attentional processes (Sadaghiani and Kleinschmidt 2016), and local representation of item content (Rohenkohl, Bosman, and Fries 2018), respectively. These observations would suggest a frequency-specific rather than a frequency-stable spatial organization of FC (Sadaghiani and Wirsich 2019).

However, computer modeling studies have shown that spatial organization of static FC in various frequency bands can be largely predicted by the structural connectivity (Cabral et al. 2014; Hansen et al. 2015; Schirner et al. 2018), suggesting some degree of frequency-invariance in the organization of FC. For example, Cabral, (2014) reported a strong correlation between empirical FC of MEG data and time series generated from theoretical computer models informed by structural connectivity. Importantly, this observation concurrently held true for multiple oscillation frequency bands suggesting that oscillation-based FC entails a frequency-independent component. Our observation of relatively strong frequency invariance agrees with this viewpoint, whilst also permitting considerably smaller frequency-specific connectivity patterns.

### Temporally independent frequency-specific FC dynamics

Our analysis of FC time-varying dynamics in each frequency band suggests that frequency-invariance likely reflects multiple frequency-specific coupling processes that enact a shared spatial organization, rather than a single broadband process. This finding may explain how frequency-specific task-evoked FC changes can co-exist with a frequency-invariant spatial organization. Although temporally independent FC dynamics in different frequency bands are consistent with temporally distinct task-evoked changes in each band, these results are dependent on the time frame considered. For instance, when assessing these dynamics over more extensive observations (i.e., longer time period or higher number of trials), the relatively unitary state- and frequency-invariant spatial organization of FC emerges. Our finding is consistent with long-term EEG and ECoG recordings that report the emergence of spatial stability only at periods >100sec (Chu et al. 2012; Kramer et al. 2011). This observation reconciles the complimentary presence of cognitive state-invariance and frequency-invariance of FC organization on the one hand and its state-responsive and frequency-specific short-term dynamics on the other.

### Phase- vs. amplitude coupling

Beyond phase coupling, this study explored the correspondence of spatial FC across cognitive states and frequency bands in the other major mode of oscillation-based connectivity: amplitude coupling. Spatial correlation values for amplitude coupling were consistently smaller than for phase coupling, but moderate to strong. The cross-state spatial correlation of both phase- and amplitude coupling was in general slightly smaller than within-resting state correlation (Fig. S4). This implies that the spatial organization of both phase- and amplitude-coupled neural connectivity is only slightly reconfigured by task manipulations. We conclude that phase- and amplitude coupling, as two different aspects of neural connectivity, are largely stable around a shared core functional architecture despite their malleable nature.

Focusing on the difference of *r* values from the within-rest condition to the cross-state conditions shown in Fig. 2 (i.e. state-dependent component of FC), we observed that task-responsiveness of the two phase- and amplitude coupling had a different profile across frequency bands (Fig. S4). The observed profiles suggest that task-related phase coupling occurs more readily in slower frequencies, while task-related amplitude-coupling enacts the higher frequencies more strongly. The dissociable task-related spatial reorganization emphasizes the distinctness of task-related phase- and amplitude coupling (Mostame and Sadaghiani 2020).

### Source leakage contributions

Could the observed state- and frequency-invariance be caused by volume conduction present in the data regardless of the cognitive state or frequency band? Although ECoG is considerably less affected by volume conduction than scalp EEG, we addressed this concern after removing zero-lag FC and distance dependence of FC. We detected a moderate reduction of the effect size of the spatial correspondence especially when suppressing zero-lag FC. Unfortunately, the conservative approach of suppressing zero-lag FC comes at the cost of removing real zero-lag connectivity whose existence (e.g., Gray et al. 1989; Rodriguez et al. 1999; Roelfsema et al. 1997) and contribution to the whole-brain connectome (e.g., Finger et al. 2016) are supported empirically and theoretically (Viriyopase et al. 2012). Importantly however, a large proportion of the data still reflected the state-invariant FC organization, indicating that volume conduction is not a primary driver of our effects. These observations emphasize the advantage of ECoG over noninvasive neurophysiological signals for investigating FC organization under minimal volume conduction effects.

### Limitations

While ECoG provides a unique window into direct intracranial recordings of the human brain, it suffers from limited electrode coverage. Thus, the observed FC organization could not be directly related to previously reported whole-brain FC networks. Importantly however, the core question regarding spatial stability of FC across cognitive states and frequency bands was not contingent upon knowing the correspondence to MRI-based networks. Moreover, the variability of electrode coverage across subjects - while it prevents comparisons across subjects-suggests that our observations are robust over different sets of brain areas.

Another limitation of ECoG is that due to its invasive nature it is only available in patients with a history of epilepsy. In particular, electrodes coverage usually includes affected brain areas to serve clinical purposes. However, data did not include electrodes and time periods with excessive inter-ictal activity. Thus, we believe that the high SNR of ECoG and its relative insensitivity to volume conduction far outweighs this potential limitation for the purposes of studying oscillation-based FC. Thereby our study adds to the wealth of prior ECoG studies successfully informing about normal brain processes (Parvizi and Kastner 2018) and FC in particular (e.g. (Betzel et al. 2019; Kucyi et al. 2018).

Another limitation ensuing from the rare opportunity of intracranial human recordings is the limited sample size. Cross-state investigations require patients with data from at least two task conditions. The overlap of *Motor* and *Rest* provided a sample size of 17 patients, which is relatively large in the context of ECoG literature. However, other tasks that we included in support of a broad representation of cognitive states were available in a subset of patients only. Nevertheless, the extraordinary SNR of intracranial recordings permits assessment of effects in individual subjects. Accordingly, the core conclusions of state- and frequency invariance were established using both group-level statistics as well as single-subject tests using subject-specific null models. The observation of all core findings in the vast majority of individual comparisons (with rigorous multiple comparisons corrections) provides strong quantitative support for the robustness of our results.

### Conclusions

Our findings suggest that spatial organization of oscillation-based FC is shared across frequency bands and is stable across a variety of tasks and rest. Invariance of oscillation-based FC across cognitive states agrees with parallel observations in the correlation structure of the aperiodic fMRI BOLD signal. This convergence suggests a universal phenomenon from the millisecond timescale of electrophysiology to the infraslow range of fMRI. These observations speak to conceptual frameworks of oscillation-based FC incorporating a largely state- and frequency-invariant spatial organization beyond state-responsive and frequency-specific short-term dynamics of FC (Sadaghiani and Wirsich 2019). This is especially important because statistical tests of task-related FC differ in the adequacy with which their assumed null distribution captures task-independent FC already present in the pre-stimulus baseline (Mostame et al. 2019). Concluding the practical implications, the state-invariant component of FC should be carefully accounted for to permit accurate observation of task-related FC changes.

## Notes

### Competing Interest Statement

The authors have declared no competing interest.

