## Supplementary materials for "Oscillation-based connectivity architecture is dominated by an intrinsic spatial organization, not cognitive state or frequency"

*Table S1 – Original number of electrodes for each subject in the dataset across rest and the other three tasks. Each row shows a subject and each column corresponds to a task. Cells containing a dash mean lack of dataset for the corresponding subject and task. For each subject, the final number of electrodes included in the analyses is equal to the minimum number in the corresponding row.*

| SUBJECT # | REST | MOTOR | SPEECH | 2BACK |
|---|---|---|---|---|
| 1 | 46 | 47 | - | - |
| 2 | 64 | 59 | - | 64 |
| 3 | 64 | 60 | - | 64 |
| 4 | 64 | 64 | - | - |
| 5 | 64 | 62 | - | - |
| 6 | 64 | 64 | - | - |
| 7 | 64 | 63 | - | - |
| 8 | 32 | 41 | - | - |
| 9 | 64 | 64 | 64 | - |
| 10 | 48 | 48 | 48 | - |
| 11 | 64 | 63 | - | - |
| 12 | 62 | 58 | - | - |
| 13 | 64 | 63 | - | - |
| 14 | 64 | 49 | - | - |
| 15 | 40 | 25 | - | 62 |
| 16 | 64 | 64 | 64 | - |
| 17 | 62 | 48 | 64 | - |
| 18 | 32 | - | - | 40 |

### 1 *Amplitude coupling*

We aimed at testing whether the state- and frequency-stable spatial organization of FC is specific to phase coupling or extends to amplitude coupling. Thus, we replicated our major findings using the amplitude coupling as the measure of FC. Results corresponding to Fig. 2, 3, and 4 of the main manuscript are shown for amplitude coupling in Fig. S1, S2, and S3, respectively. Altogether, we infer that a highly similar state- and frequency-stable spatial organization is governing both phase- and amplitude coupling.

Despite being largely stable across mental states, FC spatial patterns indeed show slight but meaningful task-related deviations from the intrinsic organization (Krienen et al., 2014). In terms of task-responsiveness, amplitude coupling showed a different profile compared to phase coupling. First, amplitude coupling shows stronger task-relevant static changes compared to phase coupling (Fig. S1 & Fig. S4; measured as the difference of within- and cross-state spatial correspondence of FC). Moreover, these task-related changes slightly increase with frequency band for amplitude coupling, while it peaks in Alpha band for phase coupling. This observation implies that low-frequency task processing is performed by both phase- and amplitude coupling changes, while high-frequency task processing is more strongly reflected in amplitude coupling changes (Fig. S4).

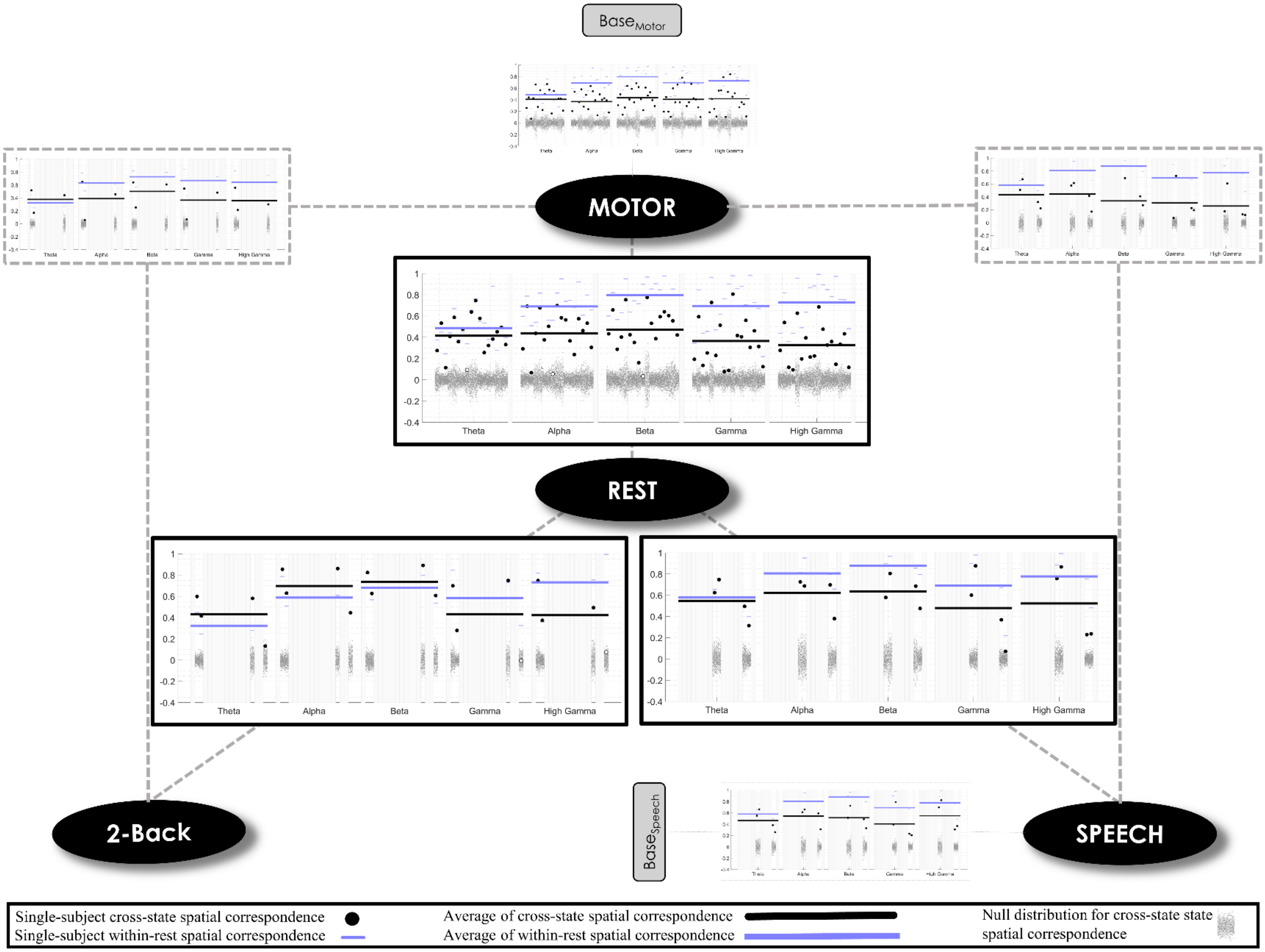


*Fig. S1 – Same as Figure 2 but for amplitude coupling (r= 0.42 ± 0.21). Black lines are substantially higher than surrogate dot clouds (gray), indicating that there is an intrinsic architecture governing the amplitude coupling mode of neural communication. Compared to phase coupling (Fig. 2), the difference between within-rest (purple) and cross-state (black) correlations were larger, implying that amplitude coupling is slightly less bound by the intrinsic architecture.*

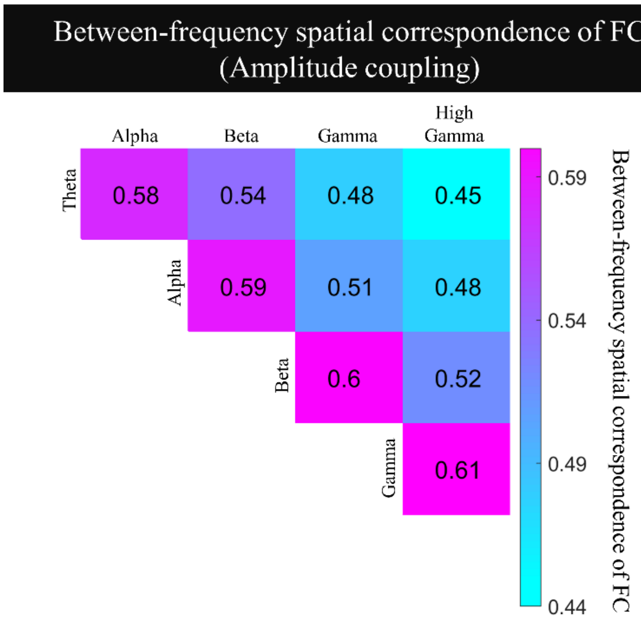


*Fig. S2 – Cross-frequency correlation of intrinsic FC for amplitude coupling. Correlation values for the amplitude coupling measure are likewise moderate to strong. Correlation values were statistically significant for all frequency pairs and individual subjects. This observation supports the hypothesis that there is a frequency-general intrinsic architecture shaping amplitude coupling paralleling the observation for phase-coupling.*

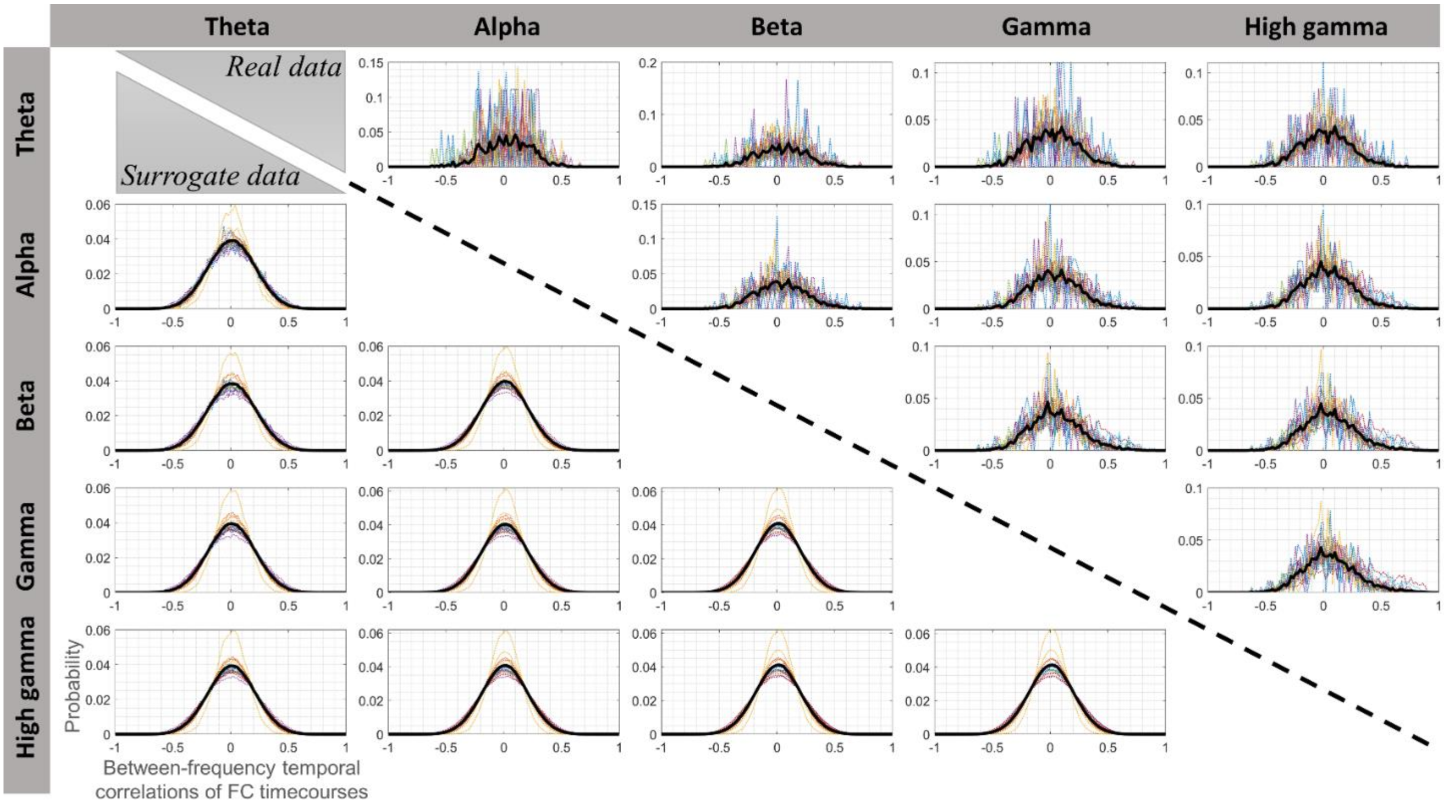


*Fig. S3 – Same as Fig. 4 but for amplitude coupling. Similarity between the histograms of the null model (lower triangle) and real data (upper triangle) shows the absence of any systematic between-frequency temporal correlation of FC dynamics. This finding suggests that that the coupling events generating the observed intrinsic FC architecture comprise several frequency-specific components with a shared spatial organization rather than a single broadband component.*

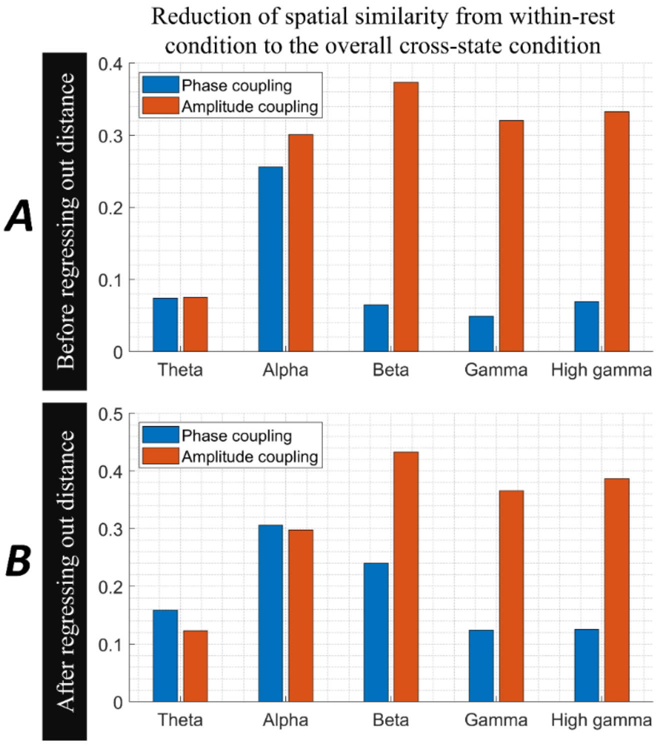


*Fig. S4 –Difference in spatial r value between the within-rest and cross-state conditions (Figures 2 & S1). Upper (or lower) panel shows the results for before (or after) regressing out the distance dependencies. In each panel, each main column corresponds to a frequency band. Blue columns correspond to phase coupling (Fig. 2), while orange columns correspond to amplitude coupling (Fig. S1). Y axis shows the r value difference between within- and cross-state conditions (within-state – cross-state). Task-related changes in FC spatial organizations - quantified by the difference of r value from within- to cross-state conditions - were overall small for both phase coupling (mean ± std= 0.10 ± 0.14) and amplitude coupling (mean ± std= 0.28 ± 0.20). Further, the impact of task for phase coupling was generally strongest in Alpha band, while it increased slightly with increasing frequency for amplitude coupling. Two-way ANOVA of frequency and coupling mode showed an interaction of* $F_{(2.49, 39.87)}$*=25.36; p=1.24e-8 (*$F_{(2.71, 43.31)}$*=25.09; p=3.85e-9 for after regressing out distance dependencies).*

#### 2 *Source leakage contributions*

We compared the cross-state spatial correspondence of FC before and after minimizing potential impact from volume conduction. We hypothesized that replicating our findings across these two conditions supports the independence of our results from volume conduction confounds. To avoid complexity, results were only replicated for one representative case of cross-state comparison (similar to one of the subplots in Fig. 2). The *Rest*-to-*Motor* comparison was selected due to the highest number of subjects. The results section of the main manuscript quantitatively elaborates on our conclusions drawn from Fig. S5.

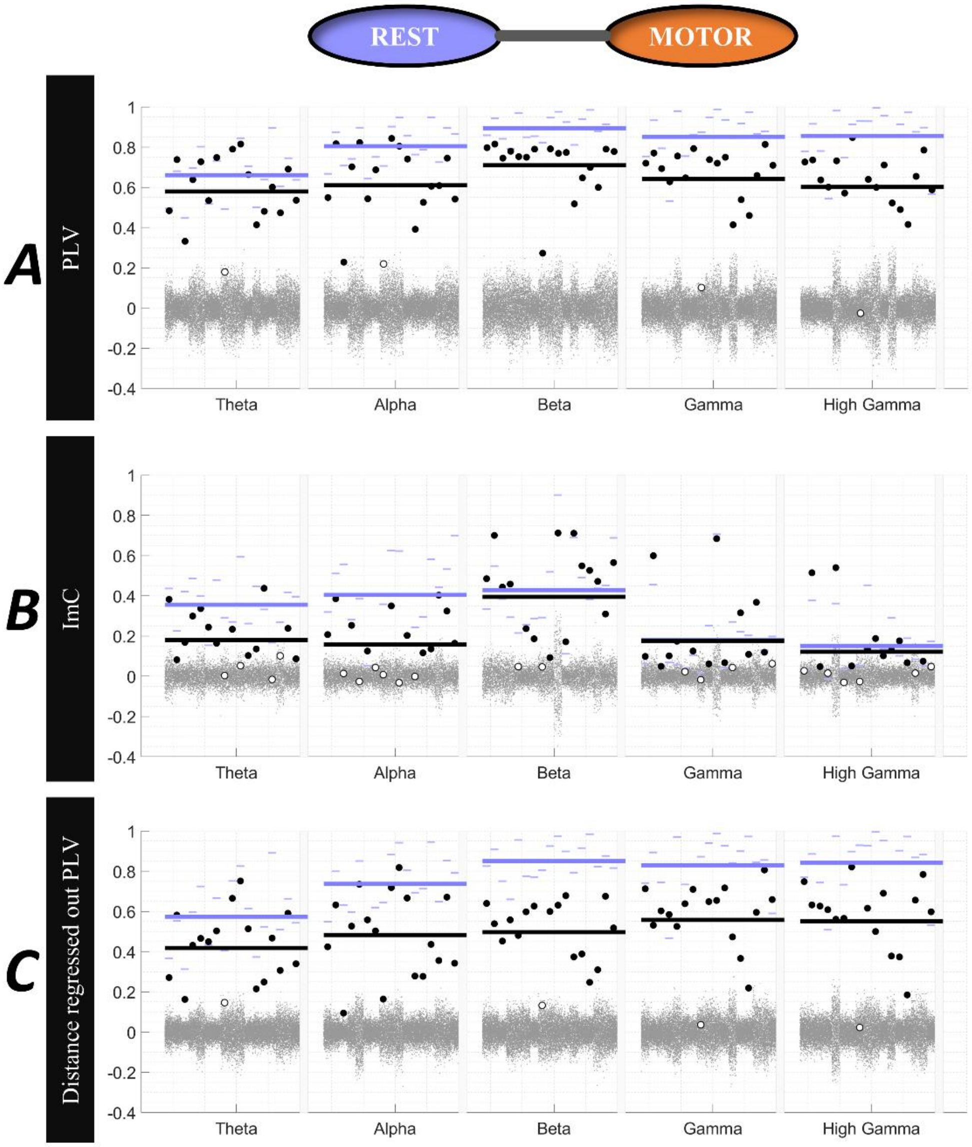


*Fig. S5 – Addressing potential contribution of volume conduction to the cross-state spatial correlation of FC. Rest-to-Motor cross-state FC correlations based on the A) PLV measure, B) ImC measure, and C) PLV measure after regressing out electrode distance. Each subplot has been plotted according to configurations of Fig. 2 in the main text. A large proportion of data shows the presence of cross-state FC correlations even after removing possible volume conduction effects, using two different approaches. This observation suggests that our major findings are not largely explained by volume conduction.*
